# Male mate choice unresolved in the mangrove rivulus

**DOI:** 10.1101/2021.02.22.432326

**Authors:** Jennifer D. Gresham, Sarah N. Bowman, Chloe M.T. Keck, Haylee M. Quertermous, Ryan L. Earley

## Abstract

Mate choice has the potential to drive phenotypic evolution because it can determine traits that increase an individual’s likelihood to reproduce (courtship behaviors, elaborate ornamentation). These traits, however, can also be detrimental for health or survival, often antagonizing the evolution of extreme phenotypes. Mangrove rivulus fish (*Kryptolebias marmoratus*) develop as self-fertilizing simultaneous hermaphrodites. Hermaphrodites overwhelmingly self-fertilize their eggs internally, but occasionally oviposit unfertilized eggs. Some individuals change sex to male after sexual maturity, essentially forgoing the reproductive assurance of selfing. In a continuing effort to understand how sex change to male is maintained this species, I designed an experiment to determine whether males act as choosers to increase their likelihood of finding unfertilized eggs for reproduction. I hypothesized that males would prefer to associate with younger hermaphrodites when given a dichotomous choice, as they lay a greater proportion of unfertilized eggs compared to older hermaphrodites. The males in this study did not show a preference for either the younger or older hermaphrodite but exhibited greater within individual variance across subtrials than among individual variation. I discuss alternative hypotheses concerning male mate choice in mangrove rivulus, which may illuminate hypotheses to be tested in this and other hermaphroditic species.

## Introduction

Mate choice occurs whenever individuals select among prospective mates, presumably using traits that inform the chooser about relative mate quality. Such choice then results in nonrandom mating. Mate choice can also occur when there is a bias in the allocation of resources (number of sperm, number of eggs, amount/quality of parental care) provided to one mate compared to other mates (Edward 2015). Selection for a variety of traits including color, size, acoustic emissions, and odor, can be attributed to mate choice, rendering it a strong evolutionary force (Andersson 1994; Edward 2015). Mate choice also drives selection for traits that may impose significant costs with regards to survival (reviewed in Jarne and Charlesworth 1993). Exaggerated traits in the chosen sex (elaborate color, larger appendages, elaborate song, courting behavior) can increase the risk of predation (Bailey 2008) and/or decrease the time that can be dedicated to acquiring food or engaging in other fitness-related activities. However, the same can be true for the chooser. Mate choice often means that the chooser must risk predation (Simcox et al. 2005), spend less time securing resources, and expend energy as they sample and assess mates (Byers et al. 2005). Ultimately, mate choice drives phenotypic evolution because it determines, along with survival and assuming the traits are heritable, which alleles will be represented in the next generation (Brooks 2002; Bleu et al. 2012), and can deepen reproductive isolation among populations and species (Panhuis et al. 2001; Roberts and Mendelson 2017; Rosenthal 2017).

The majority of mate choice studies focus on female mate choice. However, male mate choice can drive the evolution of female ornamental traits and elaborate courtship behaviors (reviewed in Schlupp 2018). Indeed, Rosenthal (2017) argues that the division of choice into “male” and “female” is too restrictive to describe the reciprocal and fluid nature of mate choice. Instead, the terms “chooser” and “courter” can apply to any sexually reproducing organism and allow for the inclusion of sequential and simultaneous hermaphroditic organisms. It is important to identify which traits of the courter are being sampled and assessed by the chooser. Courters often advertise better resources, better genes, or both before mating begins, or they can provide better post-copulatory and/or parental care after mating (Emlen and Oring 1977; Andersson 1994; Edward and Chapman 2011; Rosenthal 2017).

Mate choice in self-fertilizing simultaneous hermaphroditic species generally involves assessment of the costs and benefits of fertilizing one’s own eggs versus mating with another individual (i.e., outcrossing). When hermaphrodites can self-fertilize, their role as chooser is expanded to choosing among different mates, as well as themselves (Leonard 2006). When selfing populations are gynodioecious (hermaphrodites and females) or androdioecious (hermaphrodites and males), it is often unclear how the single-sex individuals are maintained by natural selection. These female and male courters must compete with other single-sex individuals as well as hermaphrodites that may be averse to outcrossing relative to selfing. In mobile animals, this should select for single-sex individuals that are able to find and recognize hermaphrodites, and also discriminate which individuals might be willing to outcross or identify those that are more likely to lay unfertilized eggs (Jarne and Charlesworth 1993; Martin 2007). Few studies have explored mate choice in gynodioecious or androdioecious species, although these populations offer the opportunity to test assumptions and predictions of mate choice theory against facultative sex allocation and sex change (Leonard 2006).

I explored mate choice in the androdioecious fish, the mangrove rivulus (*Kryptolebias marmoratus*, hereafter “rivulus”). Rivulus populations are comprised of selfing hermaphrodites and varying proportions of males; males result, largely, from hermaphrodites changing sex (see Harrington 1971; see also Harrington 1967 & Ellison et al. 2015 for work on the direct development of primary males). The extent of outcrossing relative to selfing increases with the proportion of males in a population (Mackiewicz et al. 2006; Tatarenkov et al. 2009). Hermaphrodites and males also are sexually dimorphic; when hermaphrodites change sex to male, they morph from having mottled brown and gray coloration to orange freckles and orange skin, usually accompanied by the loss of the caudal ocellus, an eyespot on the dorsal rim of the caudal peduncle (Harrington 1967, 1971, Soto and Noakes 1994, Scarsella et al. 2018).

Hermaphrodites fertilize their own eggs internally, and on average 93 – 95% of oviposited eggs are fertilized (see Chapter 3, Harrington 1971). Outcrossing occurs when hermaphrodites oviposit unfertilized eggs, and when males subsequently fertilize those eggs. There is no evidence that hermaphrodites outcross with each other (Furness et al. 2015). Currently, it is not known how males find unfertilized eggs oviposited by hermaphrodites. There are two possibilities of how males procure unfertilized eggs: 1) males coerce or convince hermaphrodites to oviposit unfertilized eggs in their presence, or 2) males are able to locate hermaphrodites that passively oviposit unfertilized eggs with higher frequency. Irrespective of the mechanism, males should be under strong selection to have reliable and efficient sampling algorithms to find hermaphrodites that are more willing or more likely to oviposit unfertilized eggs. I hypothesized that males would choose between hermaphrodites that may be more or less likely to lay unfertilized eggs. I presented males with two hermaphrodites of different ages and predicted that males would prefer to associate with the younger hermaphrodites. There is some histological evidence that ovarian tissue matures first (Cole and Noakes 1997), and hermaphrodites may lay unfertilized eggs before the spermatogenic tissue matures. In addition, I showed in Chapter 3 that younger hermaphrodites do not lay *only* unfertilized eggs but do lay a significantly greater proportion of their eggs unfertilized, lending additional support for this prediction.

## Methods

### Fish choice and colony conditions

Male mating preference was tested using a dual option choice arena. The behavior arena was constructed using a Sterilite^®^ plastic shoe box (item #002-02-0403). The arena was divided into three zones by external lines; a 4 cm zone around each hermaphrodite and a 5 cm neutral zone between the hermaphrodite zones. At the beginning of each trial, hermaphrodites were placed individually in a plastic breeding box that allowed the males to access both visual (unbranded, eBay, Inc. listing number 264379573516), and placed on opposite ends of the arena so that one side of each breeding box was up against the short wall of the shoebox. Placement of the hermaphrodites with respect to age and arena side was randomized for the first trial (trial A). The position of hermaphrodites was then swapped for the second trial (trial B). The focal male was placed into a 6 cm tall, 2 cm diameter, opaque white PVC pipe centered within the neutral zone. Fish were allowed to acclimate to the arena for 30 minutes before the male was released and video recording began. Fish were recorded for 30 minutes, then removed from the arena and placed back into their original containers while the arena was rinsed with 100% ethanol, then tap water, and dried. Following this break, trial B began exactly as trial A with the exception that the hermaphrodites swapped sides. To reduce external stimuli and disruptions, the arenas were separated from the rest of the room on all six sides using white lab bench paper, and the room was kept free of humans for the duration of the trial. The total number of triads (two hermaphrodites + male) was N=43.

The two hermaphrodites in each trial were chosen from the same isogenic or near-isogenic genetic lineage and generation (1 – 4 generations removed from the wild caught progenitor); they were siblings that both resulted from self-fertilization. Hermaphrodites were at least 150 days apart in age (mean ± SEM difference in age: 245 ± 10 days), but all were greater than 124 days old (mean ± SEM; old hermaphrodite age: 461 ± 14 days; young hermaphrodite age: 217 ± 9 days), well beyond the youngest age at sexual maturity (67 days, see Chapter 3). The focal male was from a genetically different lineage. I did not have any age restrictions on males (mean ± SEM: 393 ± 29 days). Each hermaphrodite pair and focal male were used in one session that was split into the two trials, A and B. All fish were kept in isolation from hatching in 25‰ water, prepared with Instant Ocean^®^ synthetic sea salt, and held in a room maintained at 26.1 ± 0.002 ºC (mean ± SEM) with 12 hour light: 12 hour dark photoperiod. Individual fish were fed a 2 ml suspension of *Artemia* nauplii daily (~1000 nauplii in 25‰ saltwater), at least one hour before the preference trials were initiated. After trial B, each fish was measured for mass (analytical balance, ± 0.0001g) and length (ImageJ software [https://imagej.nih.gov/ij/], ± 0.001mm).

### Scoring behavior videos and preference

Each video was manually scored for the number of transitions and the time spent in each zone for thirty minutes after the male was released from the central PVC pipe. The video scorers were not aware of which side contained the old and young hermaphrodite. After each video was scored, the total and proportional times in each zone were calculated, and strength of preference (SOP) scores were calculated.

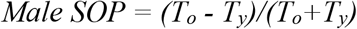

where T_o_ is the amount of time spent in the older hermaphrodite’s zone and T_y_ is the amount of time spent in the younger hermaphrodite’s zone. The SOP score can range from +1 (indicating a complete preference for the older hermaphrodite) to −1 (indicating a complete preference for the younger hermaphrodite). A SOP of zero indicates no preference. I also counted the number of times the male approached each hermaphrodite. An “approach” was counted if the male swam towards the head of the hermaphrodite to within at least one half body length.

### Statistical analysis

Normal quantile plots and model residuals distribution indicated that all variable data sets were normally distributed. To determine whether males exhibited a side bias for either the left or right zone, I used a paired t-test. To determine whether males preferred to spend more time with the older or younger hermaphrodite, I used a general linear mixed model with SOP score as the dependent variable, older hermaphrodite age, younger hermaphrodite age, number of transitions, and trial as independent fixed effects, and male ID as a random effect. I also wanted to determine whether there was a preference related to the absolute age difference of the hermaphrodites. I used a general mixed model with SOP as the dependent variable, hermaphrodite age difference, number of transitions, and trial as independent fixed effects, and male ID as a random effect. I also modeled SOP versus hermaphrodite size traits and male age as *post hoc* analyses. I used multiple general linear mixed models with SOP versus older hermaphrodite mass (or standard length), younger hermaphrodite mass (or standard length), number of transitions, and trial as independent fixed effects, and male ID as a random effect. I also used a general linear mixed model to model SOP versus the hermaphrodite mass difference plus trial, and number of transitions as independent fixed effects, and male ID as a random effect. The same model was used to determine whether standard length differences between hermaphrodites and male age were associated with SOP. Models were run in JMP Pro version 15.0.0 (“JMP^®^, Version 15 Pro” 2019).

## Results

Males exhibited a slight preference for the left zone (t_85_ = −2.1, p = 0.042); 50% of their time was spent in the left zone compared to 39.3% in the right zone. Males also did not exhibit a preference for either the older or younger hermaphrodite. Male SOP was not associated with the absolute age of either the older or younger hermaphrodite, the difference in age between the hermaphrodites, or the number of transitions that the males made among zones (Figure 4.1). My results demonstrate considerable within-male variance (considered to be a significant proportion of the residual variance), compared to among-male variance, which approached zero (random male ID variance, Table 4.1). My *post hoc* analyses tested whether male SOP might be associated with hermaphrodite size traits: mass, standard length, difference in mass, difference in standard length, and male age. My results indicate that neither hermaphrodite size nor male age were related to male SOP (Table 4.2).

**Figure 4.1.**
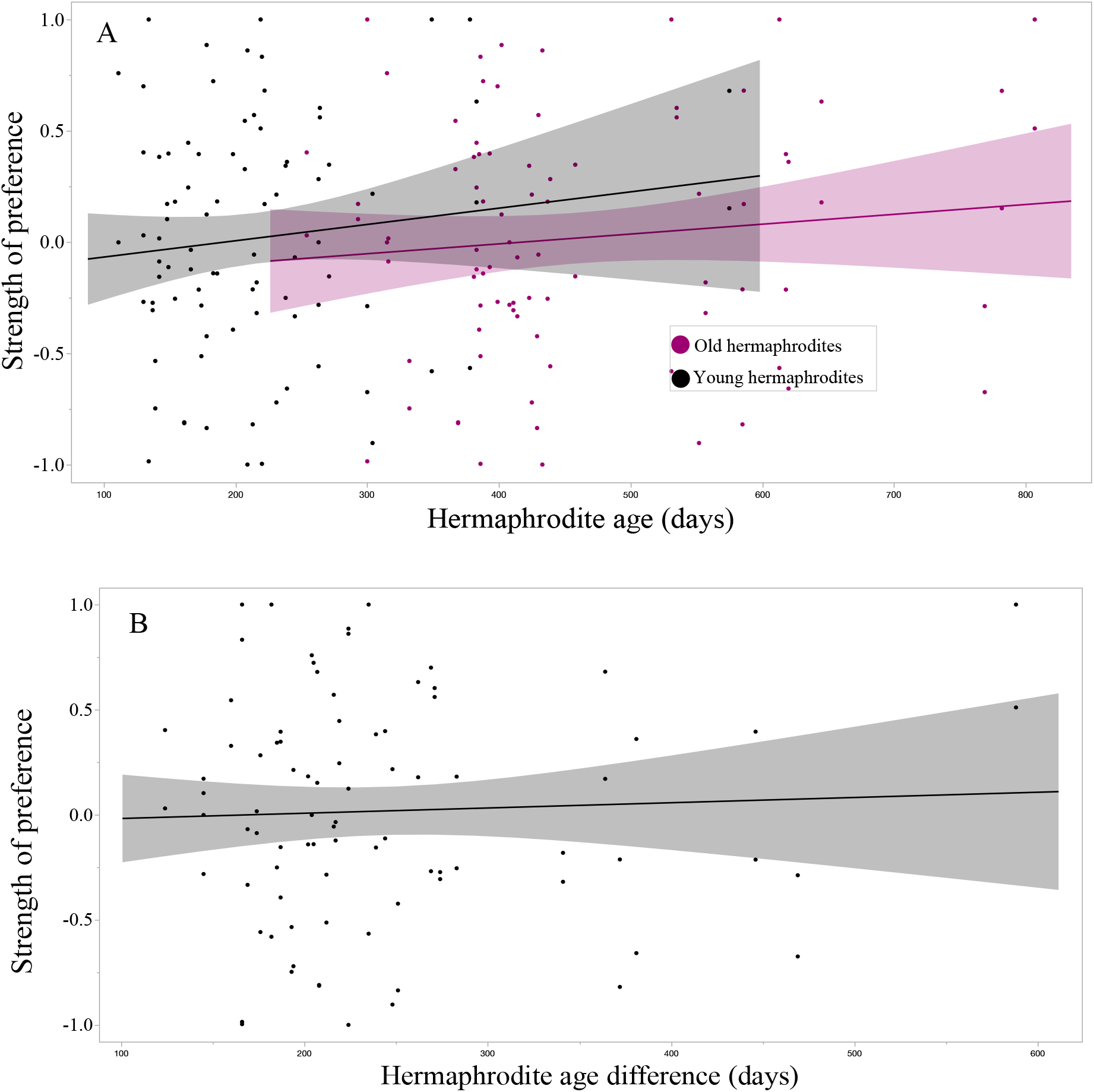
Relationships between hermaphrodite age and age difference on strength of preference. A strength of preference score of 1 indicates the male spent the entire period in the old hermaphrodite zone, while a score of −1 indicates the male spent the entire period in the young hermaphrodite zone. A) Neither the age of the older or younger hermaphrodites were significantly associated with strength of preference. B) The age difference between hermaphrodites also was not associated with strength of preference.

**Table 4.1.** Summary of models that test how hermaphrodite age, trial, and number of transitions are associated with strength of preference. F = F-ratio; df = degrees of freedom; P = P-value.

| Model effects | Estimate | F | df | P |
| --- | --- | --- | --- | --- |
| Old hermaphrodite age | 0.0002 | 0.07 | 1 | 0.80 |
| Young hermaphrodite age | 0.0006 | 0.38 | 1 | 0.54 |
| A/B trial | -0.04 | 0.38 | 1 | 0.54 |
| Number of transitions | -0.0004 | 0.27 | 1 | 0.61 |
| Hermaphrodite age difference | 0.0002 | 0.10 | 1 | 0.75 |
| A/B trial | -0.04 | 0.37 | 1 | 0.54 |
| Number of transitions | -0.0004 | 0.30 | 1 | 0.58 |

**Table 4.2.** Summary of *post hoc* analyses that test how hermaphrodite size characters, male age, trial, and number of transitions are associated with strength of preference. F = F-ratio; df = degrees of freedom; P = P-value.

| **Model effects** | **Estimate** | **F** | **df** | **P** |
| --- | --- | --- | --- | --- |
| Old hermaphrodite mass | 0.71 | 0.71 | 1 | 0.41 |
| Young hermaphrodite mass | -0.26 | 0.14 | 1 | 0.71 |
| A/B trial | -0.04 | 0.37 | 1 | 0.55 |
| Number of transitions | -0.0005 | 0.40 | 1 | 0.53 |
| Hermaphrodite mass difference | 0.43 | 0.53 | 1 | 0.47 |
| A/B trial | -0.04 | 0.36 | 1 | 0.55 |
| Number of transitions | -0.0006 | 0.46 | 1 | 0.50 |
| Old hermaphrodite standard length | -0.001 | 0.002 | 1 | 0.96 |
| Young hermaphrodite standard length | 0.004 | 0.04 | 1 | 0.85 |
| A/B trial | -0.04 | 0.34 | 1 | 0.56 |
| Number of transitions | -0.0007 | 0.57 | 1 | 0.45 |
| Hermaphrodite standard length difference | -.003 | 0.03 | 1 | 0.86 |
| A/B trial | -0.04 | 0.34 | 1 | 0.56 |
| Number of transitions | -0.0007 | 0.56 | 1 | 0.46 |
| Male age | <0.0001 | 0.07 | 1 | 0.79 |
| A/B trial | -0.04 | 0.38 | 1 | 0.54 |
| Number of transitions | -0.0005 | 0.29 | 1 | 0.59 |

## Discussion

The goal of this study was to determine whether rivulus males prefer to associate with older or younger hermaphrodites when genetic lineage and generation were controlled for. I predicted the males would prefer younger hermaphrodites, because they might be more likely to lay unfertilized eggs. Histological evidence demonstrated that ovarian tissue matures first, and implied that younger hermaphrodites might lay unfertilized eggs as “pure females” before the bifunctional gonad fully develops (Cole and Noakes 1997). In addition, I found that younger hermaphrodites lay a significantly greater proportion of unfertilized eggs (Chapter 3). The males in this study did not show a preference for either hermaphrodite based on absolute age or age difference. However, there is evidence of male preference and courtship behavior in rivulus. Male rivulus have been reported to show an association preference for hermaphrodites that were most genetically different from themselves (Ellison et al. 2013). My hermaphrodites were siblings of each other, and therefore neither was more or less genetically dissimilar from the male. There are reports that males attempt to court hermaphrodites, at least when there is only one hermaphrodite and they are not limited in their movement by a plastic box (Taylor 2012). I did not attempt to measure, nor did I observe any obvious courtship behavior, which may have been precluded by the male not being able to physically interact with the hermaphrodites. However, males exhibited very few approaches to the hermaphrodites, and some did not approach either hermaphrodite. These experiments, collectively, were conducted using a variety of environmental settings from mangrove-like mesocosms to plastic boxes. Variation in male behavior may be due to the differences in external environment as well as characteristics of the hermaphrodites, such as absolute age of the hermaphrodite or genetic similarity between the hermaphrodites and male.

My results may also suggest that males search for unfertilized eggs, not the hermaphrodites that lay them. While a few males would wait in one area of one zone for most or all of the thirty minute trial, most transitioned among the zones many times. This could be evidence that males swim around the environment searching for eggs, instead of waiting for a single hermaphrodite to lay them. In the clam shrimp *Eulimnadia texana* Packard, another selfing androdioecious species, it was concluded that males increase their potential encounters with hermaphrodites by swimming, not waiting (Medland et al. 2000). The authors also concluded that this strategy might be successful because hermaphrodites occur in high frequency, a condition observed in rivulus populations as well (Taylor 2012). Male rivulus have been shown to be less risk-averse than hermaphrodites, even when lineage is controlled (Garcia et al. 2016), supporting my observations of most males swimming around without paying much attention to the hermaphrodites. This “swim and search” strategy is also supported by the fact that hermaphrodites provide very little, if any, parental care for their eggs so hermaphrodite presence is unlikely to be a reliable cue of where unfertilized eggs can be found. In a mesocosm experiment, hermaphrodites were not observed caring for or guarding their eggs, and some were oviposited as far from the hermaphrodite’s burrow as possible (Taylor 1990). Other studies report that hermaphrodites will eat their own eggs (Harrington 1963, Wells and Wright 2017).

There are multiple reports that male fish show a preference for larger females as it presumably reflects greater fecundity (Schlupp 2018). Even in sexually dimorphic guppies (*Poecilia reticulata*) where male mate choice was thought to be insignificant, males preferred to mate with the larger of two females, and this preference was strengthened as the size difference between females increased (Dosen and Montgomerie 2004). This led us to determine whether the strength of preference of rivulus males was influenced by hermaphrodite size. Male preference was not influenced by hermaphrodite mass, hermaphrodite standard length, or mass or standard length asymmetries between the two hermaphrodites. Harrington (1963) observed that “smaller” rivulus laid fewer eggs than “larger” ones, but our lab has evidence that the opposite is true. In an experiment designed to measure phenotypic and genetic correlations between parent body size, egg size, and egg number, smaller individuals laid more eggs and larger eggs (Robertson et al., *in review*). Males do not appear to gauge the relative size of hermaphrodites or, if they do, the information appears not to be salient, and this may be further evidence that males are searching for eggs, not hermaphrodites.

How males are maintained in rivulus populations remains a mystery. There is evidence of outbreeding depression with respect to fecundity (see Chapter 1), suggesting that outcrossing and therefore males should be selected against. There also is evidence of inbreeding depression with respect to parasite load (Ellison et al. 2011) and male body size (Molloy et al. 2011), suggesting that outcrossing and males should be selected for. Further, I do not know how males find the very low proportion of oviposited eggs that are unfertilized (~6%, see Chapter 3). The lack of consistent courtship and preference behaviors reported in the few studies on rivulus makes it essential to get back to basic observational studies in the wild, preferably (at least to start) in populations where males are relatively abundant (e.g., Belizean populations). Further, whether or not hermaphrodites act as choosers and/or courters is not established in rivulus. In an experiment designed to measure association preferences, hermaphrodites preferred to spend significantly more time with males than with other hermaphrodites (Ellison et al. 2013). It is not yet clear what drives this preference, but some possibilities include outcrossing opportunities, lower risk of egg cannibalism (Wells and Wright 2017), associating with less aggressive conspecifics (Garcia et al. 2016), or minimizing resource competition for offspring. To understand precisely how androdioecy and mixed mating are maintained in rivulus, we must continue to investigate male and hermaphrodite mate choice.

